# The genome sequence of the soft-rot fungus *Penicillium purpurogenum* reveals a high gene dosage for lignocellulolytic enzymes

**DOI:** 10.1101/197368

**Authors:** Wladimir Mardones, Alex Di Genova, María Paz Cortés, Dante Travisany, Alejandro Maass, Jaime Eyzaguirre

## Abstract

The high lignocellulolytic activity displayed by the soft-rot fungus *P. purpurogenum* has made it a target for the study of novel lignocellulolytic enzymes. We have obtained a reference genome of 36.2Mb of non-redundant sequence (11,057 protein-coding genes). The 49 largest scaffolds cover 90% of the assembly, and CEGMA analysis reveals that our assembly covers most if not all all protein-coding genes. RNASeq was performed and 93.1% of the reads aligned within the assembled genome. These data, plus the independent sequencing of a set of genes of lignocellulose-degrading enzymes, validate the quality of the genome sequence. *P. purpurogenum* shows a higher number of proteins with CAZy motifs, transcription factors and transporters as compared to other sequenced *Penicillia*. These results demonstrate the great potential for lignocellulolytic activity of this fungus and the possible use of its enzymes in related industrial applications.

## 1. Introduction

Lignocellulose is by far the most abundant renewable resource on earth, and it represents a highly valuable source of raw material for different industrial processes, among them the production of second-generation biofuels such as bioethanol (Ragauskas et al. 2006). Lignocellulose contains several polysaccharides, mainly cellulose, hemicelluloses and pectin. Hydrolysis of these polysaccharides can be achieved chemically or enzymatically, the second method having the advantage of being environmentally friendly and free of potentially poisonous side-products (Blanch 2012). The saccharification process to obtain the monosaccharide components requires a variety of enzymes particularly due to the complex structure of the hemicelluloses and pectin. These enzymes are secreted by a number of bacteria and fungi (Dehority 1962; Kang et al. 2004). A thorough understanding of the lignocellulose-degrading systems is important in order to prepare enzyme cocktails that can efficiently degrade a variety of lignocellulose raw materials of different chemical composition (Rosenberg 1978). Although the study of individual enzymes (purification and characterization) may give valuable information on their properties, it is a rather time-consuming process. The recent developments in the “omics” sciences and bioinformatics have provided novel techniques for the analysis of genomes, transcriptomes and secretomes of cells and organisms, allowing a rapid identification and characterization of sets of enzymes (Conn 2003).

Fungi are the preferred sources of lignocellulolytic enzymes. Among them, the genera *Trichoderma* (Merino and Cherry 2007) and *Aspergillus* (Kang et al. 2004) have been the subject of detailed studies and several of their enzymes have been used for industrial applications. Less well known are the enzymes from *Penicillia*. However, several members of this genus have been reported to be good producers of cellulases and xylanases. A strain of *Penicillium decumbens* (Liu et al. 2013) has been used for industrial-scale cellulase production in China since 1996, and Gusakov and Sinitsyn (2012) have reviewed the production of cellulases by a set of *Pencillium* species, some of whose enzymes show superior performance to those of the better-known *Trichoderma*.

Our laboratory has used as a model for the study of lignocellulolytic enzymes a locally isolated strain of *Penicillium* (Musalem et al. 1984), which has been registered in the ATCC as MYA-38. This soft-rot fungus grows on a variety of lignocellulolytic natural carbon sources (i.e. sugar beet pulp, corncob, etc.) (Steiner et al. 1994). It secretes to the medium a large number of cellulose- and xylan-degrading enzymes, some of which have been characterized and sequenced (Chávez et al. 2006; Ravanal et al. 2010), evidencing a high and plastic lignocellulolytic enzyme activity, which may potentially have industrial applications in raw material processing.

In this article we provide a genome sequence for this *Penicillium* generated using a joint approach involving second and third generation sequencing technologies (Illumina (Bentley et al. 2008) and PacBio (Eid et al. 2009) respectively), together with annotated gene models for this fungus, analyzing novel genes focused on lignocellulolytic activity. This fungus was found, among the *Penicillia* analyzed, to have the highest number of CAZymes. An RNAseq analysis of the fungus grown on sugar beet pulp, and the independent sequencing of a set of genes of lignocellulose-degrading enzymes confirm the quality of the gene models. Due to the high number of CAZymes identified, we propose that this *Penicillium* strain is a powerful source of enzymes for the industrial lignocellulose biodegradation process.

## 2. Materials and methods

### 2.1 Fungal strain and culture conditions

*Penicillium purpurogenum* ATCC strain MYA-38 was grown in Mandel’s medium as described previously (Hidalgo et al. 1992). Liquid cultures were grown for 4 days at 28°C in an orbital shaker (200 rpm) using 1% glucose or sugar beet pulp as carbon sources. Fungus grown on glucose was used for DNA isolation and genome sequencing; cultures grown on sugar beet pulp were utilized for the transcriptome analysis.

### 2.2 Genome sequencing and assembly

For genomic DNA preparation, the fungus grown on glucose was filtered and frozen with liquid nitrogen in a mortar, and then powdered with a pestle. The genomic DNA was purified using the Genomic DNA Purification Kit (Thermo Fischer Scientific, USA) according to the manufacturer’s instructions.

Genome sequencing was carried out in two stages and with two different technologies. First, Illumina sequencing was performed on a HiSeq2000 instrument utilizing two genomic libraries: 180bp (paired-end) and 2Kb (mate-pair) insert sizes, with 100 bp read length. Second, PacBio sequencing technology was used to produce long reads (over 1Kb).

The genome assembly process was carried out in two steps. First, both Illumina libraries were assembled with ALLPAHTS-LG software (version r43019) (Gnerre et al. 2011) using a 200X coverage in order to build high quality contigs and scaffolds. Second, the Illumina assembly was scaffolded with the long PacBio reads using SSPACE-Long-Reads version 1.1 (Boetzer and Pirovano 2014).

Finally, the assembly process was validated using CEGMA (Parra et al. 2007), by aligning 248 highly conserved eukaryotic proteins to the resulting scaffolds. Since the CEGMA proteins are highly conserved, alignment methods can identify their exon-intron structures on the assembled genome, thus allowing estimation of the completeness of the assembled genome in terms of gene numbers.

### 2.3 RNASeq

The fungus was grown on sugar beet pulp as carbon source for four days and was separated from the supernatant by filtration. The mycelium was immediately processed using the TRIzol reagent (Ambion, USA) following the manufacturer’s instructions and treated with DNAse I (Invitrogen) to eliminate any remnant genomic DNA. The RNA integrity was evaluated by means of the Fragment Analyzer (Advanced Analytical Technologies, Inc., USA). The library was constructed using the Truseq Stranded mRNA LT kit (Illumina, USA), multiplexed, and sequenced in a HiScanSQ instrument (Illumina) as single reads of 101 base pairs.

Sequenced reads were mapped to the assembled genome using Tophat2 (Trapnell et al. 2009) software. Cufflinks (Trapnell et al. 2012) was used to build and quantify expression levels of RNA-Seq transcripts from aligned reads.

### 2.4 Gene discovery

For gene discovery, a variation of the pipeline described by Haas (Haas et al. 2011) was used to mark protein-coding genes in the assembled genome. This pipeline integrates evidences from different sources to create a consensus gene prediction, including ab-initio protein and transcript alignments, which are integrated with EVidenceModeler (Haas et al. 2008).

The repetitive and transposable elements of the assembled genome were identified using RepeatModeler (Price et al. 2005) and TransposonPSI (transposonpsi.sourceforge.net). RepeatModeler identifies *de novo* repeat families using kmer frequency count. These *de novo* repeats plus the latest giriRepbase (Jurka et al. 2005) were then masked using RepeatMasker (Tempel 2012).The resulting hard-masked genome was further processed using TransposonPSI which identifies transposable elements by aligning the DNA sequences to a transposon-protein database using PSI-BLAST, which were then masked using BEDtools (Quinlan and Hall 2010).

For *ab initio* gene prediction, two predictors were used: 1) AUGUSTUS (Stanke and Morgenstern (2005) (using *A. terreus* and *A. oryzae* as training sets) and 2) GeneMark-ES (Ter-Hovhannisyan et al. 2008) (self-trained). In addition, protein evidences were included using the following strategy: First, a target protein set was built by mapping the GeneMark-ES and AUGUSTUS predicted proteins against all fungal proteins from the UniProt taxonomic division using BLASTp. This set was then aligned back to the assembled genome using GenBlastG (She et al. 2011) with a cutoff e-value of 10^-10^. The resulting hits were then used as seed target proteins and mapped to the genome with GeneWise (Birney et al. 2004) in order to establish their most likely position in the genome. A match was considered evidence if the GeneWise alignment represented at least 80% of the original target protein.

The consensus gene model for each locus was produced by source-weighted integration using EVidenceModeler. Weights of 5 and 1 were assigned to GeneWise and *ab-initio* predictions, respectively. The final gene models were updated using RNAseq transcripts with PASA (Haas et al. 2003) to include splicing-isoforms, UTR tails and check the number of genes supported by RNASeq reads.

### 2.5 Functional annotation

Alignments were carried out using BLASTp version 2.2.29+ and were filtered for the first 10 hits with a cutoff e-value of 10^-5^ as suggested in the literature (Liu et al. 2013; Clarke et al. 2013). BLASTp alignments against the non-redundant RefSeq protein database (Jan 22, 2014) (Pruitt et al. 2012; Ye et al. 2006), UniProt (Jan 22, 2014) (Consortium 2014) (including Swiss-Prot, fungal taxonomic division and uniref90) and KEGG (Release 69.0, Jan 1, 2014) (Kanehisa et al. 2002) were performed to assign general protein function profiles. Protein domains were assigned using InterProScan 5.2-45.0 (Quevillon 2005) (including Pfam 27.0 (Punta et al. 2012), SUPERFAMILY 1.75 Wilson et al. 2009), SMART 7 (Letunic et al. 2012), TIGRFAMs 13.0 (Haft et al. 2013), TMHMM 2.0c (Krogh et al. 2001) PROSITE 20.99 (Sigrist et al. 2013) and PANTHER 8.1 (Mi et al. 2013) databases). Gene ontology (GO) annotations were determined using InterProScan and TransporterTP (Li et al. 2009). Enzyme Commission codes (EC numbers) and KEGG Orthology (KO) numbers were assigned using the KEGG database and BLASTp. PRI enzyme functions were assigned using PRIAM (Enzyme rel. of Mar 6, 2013) (Claudel-Renard 2003). Targeting and transmembrane signals were obtained from SignalP4.1 (Petersen et al. 2011), NetNES1.1 (la Cour et al. 2004) TargetP1.1 (Emanuelsson et al. 2000) and SecretomeP2.0 (Bendtsen et al. 2004). Transporter signals were determined using TransporterTP. Carbohydrate-activating enzymes (CAZymes) were predicted using dbCAN (Yin et al. (2012) (cutoff e-value of 10^-5^).

### 2.6 Comparative genomics and phylogenetic analysis

The annotated genes from *P. purpurogenum* were compared with twenty-one previously annotated genomes from members of the phylum *Ascomycota* (listed in Table S1) (Taxonomy IDs: 1314792, 5074, 1314791, 1314795, 1314794, 5076, 1314793, 1346256, 441960, 441959, 431241). Twenty are from the family *Trichocomaceae* (12 *Penicillium*, 5 *Aspergillus* and 3 *Talaromyces*). A cellulolytic-enzyme secreting fungus from a different family, *Trichoderma reesei*, was used as outgroup.

The assembled and annotated genomes from the *Ascomycota* reference genomes were downloaded from the JGI database (http://genome.jgi.doe.gov/). All species were assessed and compared for the number of CAZy proteins and InterPro families.

Orthologous genes between the *Ascomycota* members and *P. purpurogenum* were predicted using the OrthoMCL program (Li et al. 2003). A total of 252,277 proteins were clustered with OrthoMCL using an e-value cutoff of 10^-5^ and a moderate inflation value of 2.5 for the MCL (Enright et al. 2002) cluster method. The analysis produced 19,051 clusters, which were used to build the orthologous dendrogram. In order to compute the distance between the *Ascomycota* genomes, a membership matrix was built using orthologous clusters as rows and the genomes as columns, assigning a value of 1 if the genome belonged to the cluster and 0 if not. The final dendrogram was computed using Manhattan distances between all-pairs of genomes.

In parallel, a phylogenomics tree was constructed considering *P. purpurogenum*, the 20 aforementioned members of the Phylum *Ascomycota* and using *T. ressei* as outgroup. In order to build the phylogenomics tree, the most conserved of the orthologous clusters shared among the 22 species were selected. For each of these clusters, a multiple alignment of their corresponding amino acid sequences was performed using MUSCLE (Edgar 2004) v.3.8.3152. Groups with gaps greater than 10% of the alignment length were discarded and from this subset, only those where over 60% of their alignment length were fully or strongly conserved residues were kept, resulting in 147 clusters. Misaligned sections from these clusters sequences were removed using GBlocks 0.91b53 (Castresana 2000) and the resulting sequences were concatenated.

This concatenated sequence was used as an input for a bayesian phylogenetic analysis performed using MrBayes v 3.2 (Huelsenbeck and Ronquist 2001). The analysis was carried out partitioning by gene and applying the Jones model to each partition, as it was the one with the highest posterior probability (1.0) with parameters unlinked across partitions. Two independent runs of 1.000.000 generations were conducted with tree sampling every 1.000 generations. A consensus tree was built from the two runs using a burn-in value of 25%. Finally, FastTree (Price et al. 2009) was used to produce another phylogenomics tree using the aforementioned concatenated sequence to confirm MrBayes results by a different approach. FastTree was run with default parameters for amino acid sequence.

In addition, for a set of members of the *Penicillium, Aspergillus* and *Talaromyces* genera, sequences were downloaded for marker genes BenA, CaMand RPB2 (β-tubulin, calmodulin and RNA polymerase II second largest subunit, respectively) (Table S2). This set of sequences was used to perform a Blast search against *P. purpurogenum* genes.

### 2.7 Manual annotation of CAZymes

dBCAN searches for genes coding for glycosyl hydrolases (GH), glycosyl transferases (GT), polysaccharide lyases (PL) and carbohydrate esterases (CE) from the CAZy database were performed. From the results obtained (Table S3), all CAZy genes with an e-value of less than 10^-40^ were curated with the help of Apollo (Lewis et al. (2002). In addition, each entry was subjected to BLASTp and CD-search (Manchler-Bauer and Bryant 2004) to manually confirm or improve the automatic annotation.

### 2.8 Accession numbers

Genome and RNASeq sequences were submitted to the Sequence Read Archive (SRA) and can be downloaded from BioProject SRP055745. SRA numbers for Illumina-overlapping, Illumina-jumping, PacBio and RNASeq reads are SRR1823665, SRR1823673, SRR1823957 and SRR1824012 respectively. Assembled genome and related annotation files can be downloaded from http://ppurdb.cmm.uchile.cl/material.

## 3. Results and discussion

### 3.1 Genome sequencing and assembly

The Illumina sequencing process produced 221.7 million reads, representing a raw coverage of 568X. The PacBio sequencing produced 975,179 reads larger than 1kb, representing a raw coverage of 44.25X, with an N50 of 1,771 base pairs.

The assembled genome has 158 scaffolds with a length of 36.2 Mb (Table 1); 90 percent of the genome length is covered by only 49 scaffolds, indicating that the assembly is highly continuous. The assembly length covers 92% of the estimated genome size obtained from the kmer spectrum analysis of 39.2 Mb at a K = 25 scale (Figure S1). This estimated genome size, however, is not consistent with a previously reported genome length (Chávez et al. 2001), which was obtained with contour-clamped homogeneous electric field gel electrophoresis, resulting in an estimated genome size of 21.2 Mb. We attribute this discrepancy to a lack of resolution of the larger chromosomes by pulse field electrophoresis.

**Table 1.** Assembly summary.

|  | Illumina<br>Assembly | PacBio +<br>Illumina |
| --- | --- | --- |
| Number of contigs | 836 | 687 |
| Total contigs size (bp) | 35,743,205 | 35,951,513 |
| Min contig length (bp) | 1,000 | 947 |
| Max contig length (bp) | 462,380 | 665,523 |
| N50 contigs (kb) | 131 | 230 |
| Number of scaffolds | 582 | 158 |
| Total scaffolds size (bp) | 35,858,128 | 36,207,399 |
| Min scaffolds length (bp) | 1,000 | 977 |
| Max scaffolds length (bp) | 1,235,734 | 2,800,880 |
| N50 scaffolds (kb) | 213.4 | 838.183 |
| N90 scaffolds (kb) | 12.2 (249) | 188.1 (49) |

The N90 and N50 scaffolds are the length of the scaffolds covering 90 and 50 percent of the genome, respectively. The N90 and N50 were calculated using 39 Mb as genome size. In parenthesis are the numbers of contigs/scaffolds required to cover 90% of the genome length

The scaffolding with PacBio reads incremented the scaffold N50 fourfold in relation to the high-coverage using only Illumina reads (Table 1). The genome was assembled with the highest sequence coverage and it is, to our knowledge, the first *Penicillium* assembled with PacBio technology.

CEGMA analyses (Table S1) indicate that 99.6% of the 248 ultra-conserved core eukaryotic genes are present in the assembled genome, and 98.4% of them were considered complete; in addition, our CEGMA numbers are in agreement with respect to the reference fungal genomes (Table S1). Further, 93.1% of the RNASeq reads aligned to the assembled genome. Collectively, these results indicate that our assembled genome is highly continuous and captures the majority, if not all, the protein-coding genes present in the genome.

### 3.2 Repeat masking and gene discovery

Sixty-five families of repetitive elements were *ab-initio* found and classified with RepeatModeller. 22%, 14%, 9% and 55% were classified as Long Terminal Repeats (LTR), DNA transposons, Long Interspersed Elements (LINE) and unknown elements, respectively. RepeatModeller families in conjunction with Repbase (Feb 3, 2014) masked 9.83% (3.8 Mb) of the assembled genome. The major family corresponds to LTR elements, specifically to the Gypsy family, which represents 51% of the masked sequences. Similar results have been reported previously for fungal genomes (Muszewska et al. 2011).

One hundred and eleven transposable elements were identified which masked another 0.2% (76 Kb) of the genome. 3.8 Mb were thus masked, representing a 10% of the assembled genome, which is consistent with the estimated repeat percentage from the kmer analysis.

The *ab-initio* gene predictors rendered 33,544 gene models. A total of 1,642,820 non-redundant consolidated proteins from UniProt were mapped against our gene models, identifying a total of 10,811 unique homologous proteins. This set gave 16,764 hits against the whole genome with GeneBlast. 12,559 gene models were obtained with GeneWise using proteins as evidence. Using EVidenceModeler and PASA, 11,057 consolidated genes were found (Table 2). These predicted gene sequences account for 59% of the *P. purpurogenum* genome, with an average gene length of 2,105 bp. On average, each gene contains 3.3 exons and 2.2 introns (Table 2). Comparison of gene models between the chosen fungal genomes (Table S1) shows that our genome has on average larger genes and transcripts. This is explained by the annotation of UTR-tails with PASA and the absence of UTR-tails annotations on the reference fungal genomes. No significant difference on the average protein length, number of exons and average intron length is observed when comparing genomes, supporting the aforementioned observation.

**Table 2.** Relevant statistics of gene discovery.

| Attribute | Stats |
| --- | --- |
| Number of genes | 11,057 |
| Number of mRNAs | 11,555 |
| Number of exons | 37,145 |
| Number of coding DNA sequences | 35,980 |
| Number of introns | 24,653 |
| Exons per gene | 3.3 |
| Coding DNA sequences per gene | 3.2 |
| Introns per gene | 2.2 |
| Genes alternative spliced | 454 |
| Percentage of genes alternative spliced (%) | 4.1 |
| Average gene length(bp) | 2,105 |
| Average exon length (bp) | 612 |
| Average intron length (bp) | 92 |
| Average distance between genes (bp) | 1,349 |
| Genome coding(%) | 59 |
| Number of tRNAs | 170 |

A consensus gene set was built with EVM and PASA using evidence from *ab-* *initio*, protein and transcript alignments

A total of 9,585 genes (87%) were confirmed by at least 10 reads from RNASeq data, indicating that our gene models are well supported and in agreement with transcriptome data.

Out of the 11,057 genes predicted, 94% (10,422) were successfully annotated (Table S4) using standard functional protein databases. We were able to assign InterPro Number, GO Numbers and CAZy IDs to 80%, 63% and 8% of the gene models (Table S4), respectively.

### 3.3 Further validation of the genome sequence

In addition to RNASeq data, the quality of the genome sequence can be validated by the result of independent sequencing of individual genes. Table 3 lists a set of lignocellulose-degrading enzymes which have been studied and sequenced either prior to the availability of the draft genome or as a result of mining the genome. In all cases, the gene sequences obtained utilizing the Sanger method agree with the sequence predicted by the genome. These studies, which show properties of novel enzymes, are also proof that *P. purpurogenum* is a powerful source for the discovery and analysis of new lignocellulolytic enzymes with potential biotechnological applications.

**Table 3.** Characterized lignocellulose-degrading enzymes from *P. purpurogenum* whose gene sequence was determined independently

| Enzyme | CAZy family | Gene ID | GenBank | Reference |
| --- | --- | --- | --- | --- |
| Endoxylanase A | GH 10 | PPSCF00028.25 | AAF71268 | Chávez et al. 2001 |
| Acetyl xylan esterase II | CE 5 | PPSCF00020.380 | AAC39371 | Gutiérrez et al. 1998 |
| Arabinofuranosidase 1 | GH 54 | PPSCF00061.89 | AAK51551 | Carvallo et al. 2003 |
| Arabinofuranosidase 2 | GH 51 | PPSCF00010.19 | EF490448 | Friz et al. 2008 |
| Arabinofuranosidase 3 | GH 43 | PPSCF0002.743 | FJ906695 | Ravanal et al. 2010 |
| Arabinofuranosidase 4* | GH 54 | PPSCF00024.186 | AGR66205 | Ravanal and Eyzaguirre 2015 |
| Pectin lyase | PL 1 | PPSCF00015.779 | KC751539 | Pérez-Fuentes et al. 2014 |
| Exo-arabinanase* | GH 93 | PPSCF00035.45 | KP313779 | Mardones et al. 2015 |

\*The gene was mined from the genome and later sequenced

Manually annotated CAZyme genes present in the *P. purpurogenum* genome In order to obtain a more precise identification of CAZy genes and their enzymes, a manual annotation of a sub-set of the CAZymes (e value of 10^-40^) from those originally assigned by dbCAN with an e-value of 10^-5^ was performed. The result is presented in Table S5. A total of 306 genes have been manually annotated, coding for 219 glycoside hydrolases (GH), 49 glycoside transferases (GT), 6 polysaccharide lyases (PL) and 31 carbohydrate esterases (CE), thus confirming the high potential for lignocelluloses degrading enzymes present in the fungus..

### 3.4 Comparative functional analysis

In order to identify potentially unique novel functions for *P. purpurogenum*, we constructed a tree map of GO-terms using 688 genes without orthologs among the selected *Ascomycota* species arranged according to their uniqueness (Supek et al. 2011). Figure S2 shows a uniqueness for genes related to glycolipid transport, Krebs cycle, proteolysis, metabolism of L-arabinose and stress response.

In Figure S3 we compare the functional profile of *Ascomycota* genomes using InterPro domains. We observe that *P. purpurogenum* has, on average, lower number of genes related to specific central metabolic processes such as protein kinase signals, methyltransferases, oxidoreductases and alpha/beta hydrolases. In addition, it potentially has a lower capacity to incorporate amino acids into the cell and has a lower number of genes associated to Zinc and homeodomain transcription factors. However, we observe in *P. purpurogenum* a higher capacity in terms of gene dosage for functions related to pectin lyase, major facilitator superfamily (MFS) and glycoside hydrolases (involved in plant cell-wall degradation), general substrate transport, gene regulation and degradation of biomass. The overrepresented terms suggest that *P. purpurogenum* is more specialized in the transport of elements across the cellular membrane (IPR016196 and IPR005828). This is a signal that the fungus has a stronger performance in the secretion of extracellular proteins and import of extracellular sugars. In addition, it is interesting that *P. purpurogenum* has an elevated number of proteins with IPR011050 domain, a domain associated to pectin lyases; this is consistent with the good growth of the fungus when cultivated on rich pectin carbon sources such as sugar beet pulp.

The differences in the number of genes related to transcription factors, pectin lyases and glycoside hydrolases, with respect to the average found in *Ascomycota* genomes, suggest that *P. purpurogenum* is an interesting model to explore novel mechanics for biodegradation of lignocellulose and its regulation.

To provide further insight into the aforementioned capacity of *P. purpurogenum* for biodegradation of lignocellulosic compounds, we compared the *Ascomycota* genomes using the Carbohydrate-ActiveEnZymes classification system. The CAZy family diversity per genome is shown in Figure S4 (the raw data of putative predicted CAZYmes utilized to construct Figure S4 have not been included in the article due to their excessive length. They are available from the authors upon request). There is no great difference in terms of CAZy family numbers among the *Ascomycota* genomes as shown in Figure S5 (a large core of 129 CAZy families is shared by all *Ascomycota* genomes). However, Figure S6 shows that the number of genes related to each CAZy family varies among the genomes. Thus, we may conclude that the performance for biodegradation of lignocellulose may be more related to gene dosage than to differences in family diversity, and in this respect *P. purpurogenum* presents a higher potential than the other fungi analyzed.

### 3.5 Comparative phylogenetic analysis

CEGMA analysis shows that all *Ascomycota* genomes in Table S1 are complete in terms of gene models, thus allowing comparative genomic analysis at the gene level without the bias produced by incomplete genomes.

Orthologous analysis whit OrthoMCL produced 19,052 ortholog groups. To represent pairs distances among *Ascomycota* genomes in terms of ortholog clusters, we built an orthologous dendrogram (Figure S7) based on Manhattan distances (See Materials and methods). The orthologous dendrogram locates *P. purpurogenum* close to other *Penicillia* and far from *Talaromyces* in term of shared orthologous clusters (on average 5,502 clusters of distance to the *Talaromyces* species).

To confirm the previous observation we devised a phylogenetic analysis using a core of 147 conserved clusters among *Ascomycota* genomes (Figure 1), using *Trichoderma reesei* as outgroup and the super-matrix phylogenetic approach (See Materials and methods).

**Figure 1.**
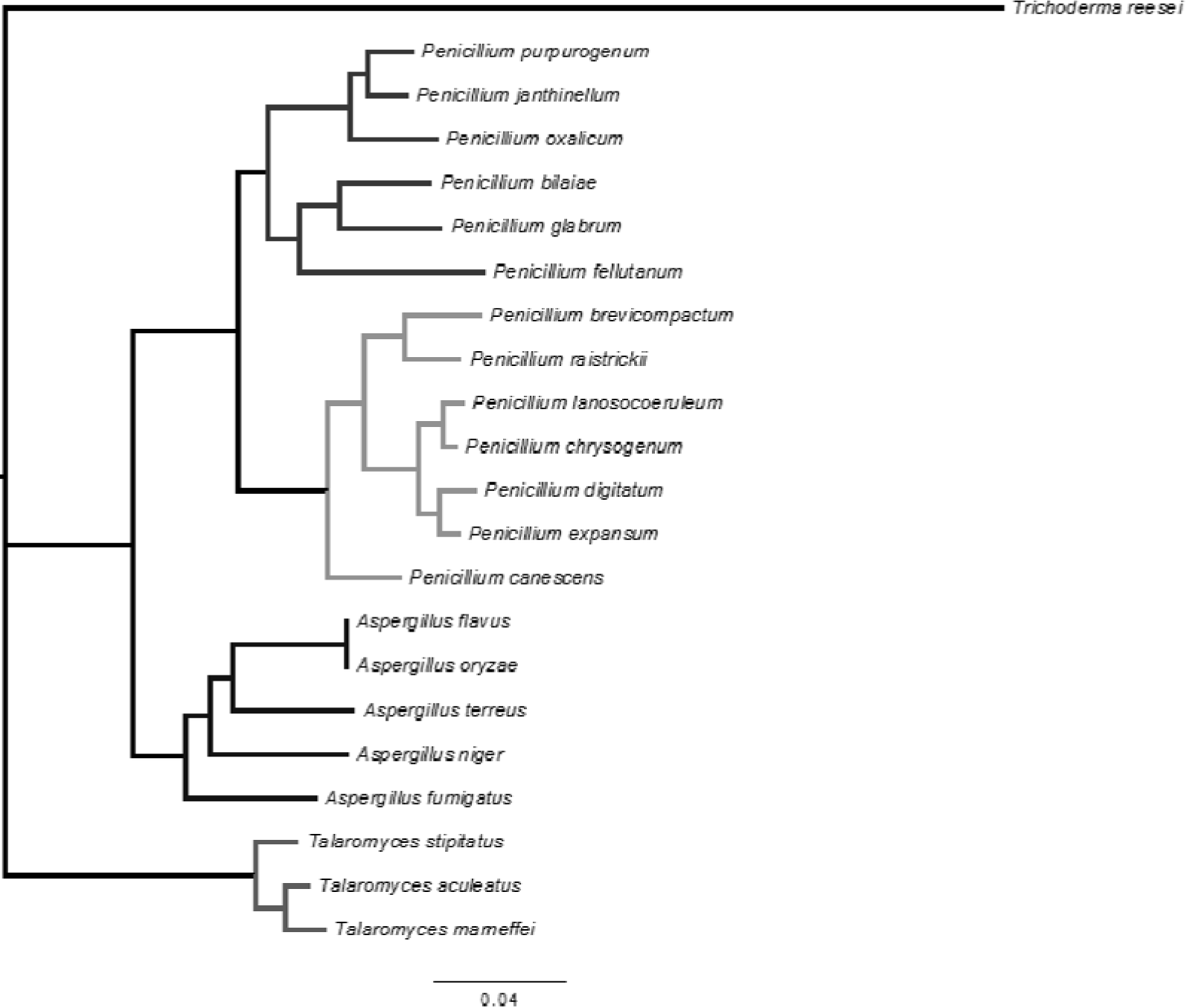
Phylogenetic tree of the Phylum *Ascomycota* inferred using mrBayes and FasTree analysis, considering the deduced amino acid sequences of 147 conserved genes. *T. ressei* was used as outgroup.

Our phylogenetics analysis (Figure 1) locates *P. purpurogenum* in the *Penicillium* genus and recovers the known phylogenetic relations among *Trichoderma, Aspergillus*, *Talaromyces* and *Penicillium* (Samson et al. 2011; Houbraken et al. 2014) where *Penicillium* is located closer to *Aspergillus* than to *Talaromyces*. Also, within *Penicillium*, we recover the current two subgenus classification (Houbraken et al. 2014), which corresponds to *Aspergilloides* (*Penicillium* A) and *Penicillium* (Penicillium B) and we locate *P. purpurogenum* as member of the subgenus *Aspergilloides*.

In addition, we compiled a set of sequences for genes BenA, CamA and RPB2 from several species of the genera *Penicillium* (212 sequences), *Aspergillus* (227 sequences) and *Talaromyces* (68 sequences) (Table S2). It has been previously suggested that these genes are well suited as markers for the identification of *Penicillium* species (Visagie et al. 2014). We found that for these three markers, *P. purpurogenum* MYA-38 had the highest identity with sequences from *Penicillium* species, although no exact match was found. The closest match was with *P. ochrochloron* with an average identity of 98.6%. These findings support the classification of our fungus as a *Penicillium*.

## 4. Conclusions

Combining the PacBio and Illumina technologies we have succeeded in producing a high quality genome sequence for *P. purpurogenum*.

The genome annotation of our strain reveals a higher secretion activity as compared to other fungi analyzed, as well as an important number of CAZy and transporter proteins, indicating a high potential for secreting enzymes involved in lignocellulose biodegradation.

The comparative genomic analysis shows that all *Ascomycota* genomes have a similar repertory and share a large core of CAZyme families. However, they differ in gene dosage, suggesting that this may relate to a different performance in lignocellulose biodegradation.

The availability of a high-quality genome sequence presents a platform for the searching of genes coding for enzymes of potentially novel functions. In addition, this genome is a valuable tool for the analysis and understanding of other fungal genomes.

## Acknowledgements

The authors thank Dr.Shahina Maqbool (Albert Einstein College of Medicine of Yeshiva University, New York, USA) for performing the Illumina DNA sequencing; Dr. Rodrigo Gutiérrez and Dr. Tatiana Kraiser (Pontificia Universidad Católica de Chile, Santiago, Chile) for the RNASeq sequence and Dr. Olivier Fedrigo (Duke University, USA) for the PacBio sequencing.

## Supplementary material

Supplementary data to this article (Tables S1 to S5; Figures S1 to S7) can be found online at

